# Quantitative Tyr Phosphoproteomic Analyses Identify Cholesterol as a Master Regulator of the Tumor Adaptive Resistance to MAPK Inhibition

**DOI:** 10.1101/2021.05.20.444617

**Authors:** Xu-Dong Wang, Chiho Kim, Yajie Zhang, Smita Rindhe, Melanie H. Cobb, Yonghao Yu

## Abstract

Although targeted inhibition of the MAPK pathway has achieved remarkable patient responses in many cancers with MAPK hyperactivation, the development of resistance has remained a critical challenge. Besides genomic resistance mechanisms, adaptive tumor response also underlies the resistance to targeted MAPK inhibitors. It is being increasingly appreciated that such bypass mechanisms often lead to the activation of many pro-survival kinases, which complicates the rational design of combination therapies. Here we performed global tyrosine phosphoproteomic (pTyr) analyses and demonstrated that targeted inhibition of MAPK signaling in melanoma cells leads to a profound remodeling of the pTyr proteome. Intriguingly, many of these kinases contain a cholesterol binding motif, suggesting that altered cholesterol metabolism might drive, in a coordinated fashion, the activation of these kinases. Indeed, we found a dramatic accumulation of intracellular cholesterol in melanoma cells (with BRAF^V600E^ mutations) and non-small cell lung cancer cells (with KRAS^G12C^ mutations) treated with MAPK and KRAS^G12C^ inhibitors, respectively. Importantly, depletion of cholesterol not only prevented the MAPK inhibition-induced feedback activation of pTyr singling but also enhanced the cytotoxic effects of MAPK inhibitors, both in vitro and in vivo. Taken together, our findings provide the evidence suggesting that cholesterol functions as a master regulator of the tumor adaptive response to targeted MAPK inhibitors. These results also suggest that MAPK inhibitors could be combined with cholesterol-lowering agents to achieve a more complete and durable response in tumors with hyperactive MAPK signaling.

## Introduction

Aberrant activation of the Ras-BRAF-MEK-ERK pathway underlies the pathogenesis of many human malignancies. Genetic mutation events (e.g., *BRAF/Nras* mutations in melanoma and *Kras* mutations in non-small cell lung cancer) result in the constitutive activation of this mitogen-activated protein kinase (MAPK) signaling pathway, which leads to uncontrolled cell proliferation, evasion of cell death, and eventually, oncogenic transformation. The identification of these genetic alterations provides the strong rationale to target these unique tumor-acquired vulnerabilities, with multiple MAPK pathway inhibitors recently approved by the FDA (e.g., BRAF and MEK inhibitors for the treatment of melanoma). Furthermore, MAPK inhibitors (e.g., KRAS^G12C^ inhibitors) are evaluated, either alone as single agents or in combination with chemo- and radiation-therapies, against a wide variety of other solid tumors.

Although many cancer patients with these actionable mutations initially respond to the targeted therapies, single agent MAPK inhibitors rarely achieve a durable response, and therapeutic resistance almost invariably occurs. Although the underlying mechanisms are complex, a universal theme of the resistant phenotype is the sustained pro-survival signaling in the cells that evade these MAPK inhibitors (1, 2). This is exemplified by the identification of the “acquired resistance” mechanisms, which includes the development of activating MEK mutations (e.g., MEK1^C121S^ (3) and MEK2^Q60P^(4)), or the expression of alterative splicing isoforms of BRAF (e.g., p61BRAF^V600E^)(5) in relapsed melanoma patients. In addition to genomic resistance mechanisms, an emerging resistance mechanism to MAPK inhibitors involves the reshaping of the signaling network in tumor cells that allows these cells to adapt to the inhibition of this key survival pathway (termed as “adaptive response”) (6, 7). Indeed, increased activation of receptor tyrosine kinases (RTKs), including the platelet-derived growth factor receptor (PDGFR)-β(8) and insulin-like growth factor-I receptor (IGF-IR)(9) is frequently observed in post-treatment biopsy samples obtained from melanoma patients. Enhanced phosphotyrosine (p-Tyr) signaling then results in the (re)activation of downstream pathways (e.g., MAPK and the phosphatidylinositol-3-OH kinase (PI(3)K)-AKT), and thereby provides an alternative means to sustain tumor growth under the MAPK-inhibited conditions. Besides in the context of MAPK-inhibited melanoma, the activation of “bypass” signaling mechanisms is a nearly universal paradigm for targeted therapies in multiple cancer types (10), including the activation of EGFR in KRAS^G12C^ inhibitor-treated non-small cell lung cancer (NSCLC) cells (11), IGF1R in mTORC1 inhibitor-treated breast cancer cells (12) and FGFR in Met inhibitor-treated leukemia cells (13)

More recent studies have shown that a common feature of the adaptive tumor response to targeted therapies is often not the activation of one or two kinases, but rather a systematic remodeling of the receptor tyrosine kinome (14, 15). Furthermore, different sets of kinases can be activated, in a context-specific manner, in various tumor cells following pharmacological perturbations (16). Because of the simultaneous engagement of multiple, and often redundant survival signals, combination therapies involving a targeted therapeutic agent and an additional kinase inhibitor are less likely to succeed.

A key question is then, upon the treatment of targeted therapies, whether there exists a master regulator that controls the concerted activation of multiple RTKs? We envision that a thorough understanding of the molecular nature of such a master regulator will greatly facilitate the design and implementation of improved therapeutic strategies to achieve more complete and sustained responses in a broad spectrum of solid tumors. Here, using high sensitivity mass spectrometry, we performed unbiased assessment of the global p-Tyr proteome in melanoma cells treated with various targeted MAPK inhibitors. The results showed that MEK/BRAF inhibitor treatment results in a profound remodeling of the tyrosine kinome, leading to the widespread activation of p-Tyr-mediated signaling events. Intriguingly, subsequent bioinformatic interrogation of our quantitative proteomic dataset pointed to a common cholesterol-binding motif among the activated Tyr kinases, suggesting that altered cholesterol metabolism might be linked to the activation of bypass p-Tyr signaling. Indeed, dramatic accumulation of cholesterol was observed in BRAF^V600E^ melanoma and KRAS^G12C^ lung cancer cells treated with various MAPK inhibitors. Depletion of cholesterol completely abrogated the activation of p-Tyr signaling in melanoma cells treated with BRAF/MEK inhibitors. Finally, we showed that a combination therapy involving MAPK inhibitors and cholesterol synthesis blockers overcome adaptive resistance, leading to greatly improved cytotoxicity in both in vitro and in vivo models of melanoma and NSCLC. Our findings thus point to cholesterol as a master regulator of the feedback signaling program, which warrants the evaluation of the proposed combination therapy in future clinical studies.

## Results

### Quantitative Tyr phosphoproteomic analyses of the adaptive response in melanoma cells

As a model system, we used two melanoma cell lines, A375 and Mel-Juso, to investigate the feedback regulation of p-Tyr signaling by the MAPK inhibitors. These two cell lines have activating mutations (i.e., BRAF^V600E^ in A375 and NRAS^Q61L^ in Mel-Juso), and therefore are characterized by their hyperactive MAPK signaling. Immunoblotting analyses confirmed that the acute treatment of A375 cells with a BRAF inhibitor PLX4032 (Vemurafenib) abrogated the phosphorylation of its key downstream target proteins, including MEK, ERK, p90 ribosomal S6 kinase (RSK) and S6 (Fig. S1A, S1B, Figure S1-source data 1). Consistent with previous reports, we observed the reactivation of these proteins after prolonged PLX4032 treatment (Fig. S1B, Figure S1-source data 1). Similar findings were obtained when these cells were treated with a MEK inhibitor (AZD6244, Selumetinib) (Fig. S1C, Figure S1-source data 1). In Mel-Juso cells (BRAF^WT^/NRAS^mut^), AZD6244 caused the adaptive resistance whereas PLX4032 induced transactivation of RAF dimers, leading to the rapid and paradoxical activation of MAPK signaling (Fig. S1D and S1E, Figure S1-source data 1) (17).

To characterize the feedback regulation of p-Tyr signaling, we performed four sets of large-scale, SILAC (stable isotope labeling by amino acids in cell culture)-based quantitative tyrosine phosphoproteomic experiments. In the first experiment (A375_PLX4032), we treated the light and heavy A375 cells with DMSO and PLX4032, respectively, for 48 hrs (Fig. 1A and Fig. S1A, Figure S1-source data 1). Cells were harvested, and the lysates were combined at a 1:1 ratio. Proteins were digested, with the resulting peptides subject to p-Tyr peptide enrichment and quantitative LC-MS/MS analyses (see Materials and Methods text for the detailed description of the experiment). Similar experiments were also performed on A375 cells treated with/without AZD6244 (A375_AZD6244), Mel-Juso cells treated with/without AZD6244 (Mel-Juso_AZD6244) and Mel-Juso cells treated with/without PLX4032 (Mel-Juso_PLX4032).

**Fig. 1.**
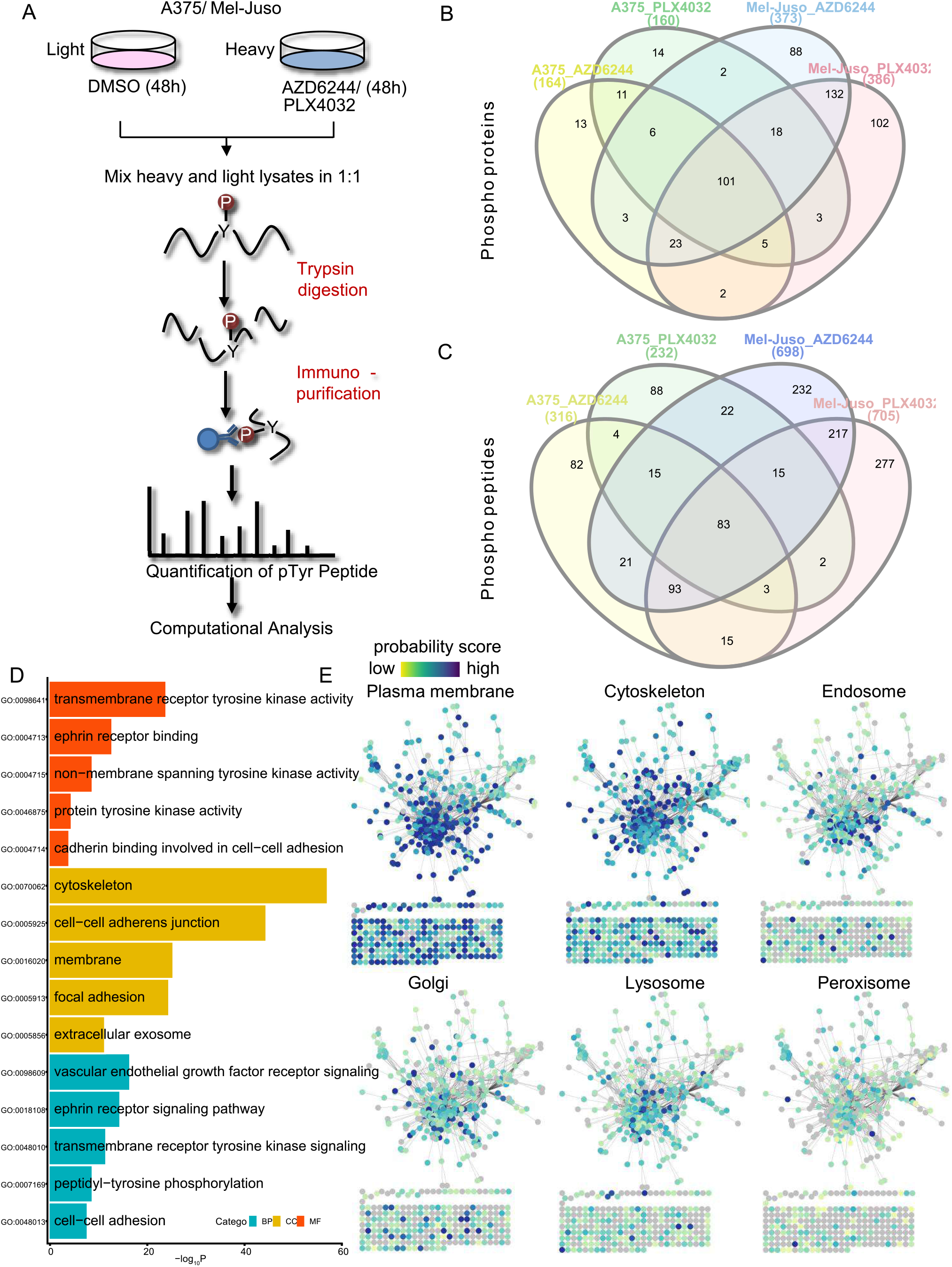
(A) The general schematic of the quantitative Tyrosine phosphoproteomics workflow. Briefly the SILAC labeled A375 and Mel-Juso cells were treated with AZD6244 or PLX4032 for 48 hours as indicated, followed by rapid lysis, protein extraction, and digestion into tryptic peptides. Then the peptides were enriched by the anti-Phospho-Tyr mAb beads. The resulting phosphopeptides were analyzed by LC-MS/MS experiments. (B) Venn diagrams showing the overlap between the phosphorylated proteins identified in A375_AZD6244, A375_PLX4032, Mel-Juso_AZD6244 and Mel-Juso_PLX4032. (C) Venn diagrams showing the overlap between the phosphorylated peptides identified in A375_AZD6244, A375_PLX4032, Mel-Juso_AZD6244 and Mel-Juso_PLX4032. (D) Gene Ontology (GO) analysis of the phosphorylated proteins identified across the four sets of experiments. The most represented GO terms related to Biological Process (BP), Cellular Component (CC) and Molecular Function (MF) are shown (Benjamini-Hochberg corrected *P* value). (E) The COMPARTMENTS resource based subcellular localization analyses of the phosphorylated proteins identified across the four sets of experiments. The color of each protein represents the confidence scores.

From these four sets of SILAC experiments, we were able to identify a total of 316 (A375_AZD6244), 232 (A375_PLX4032), 698 (Mel-Juso_AZD6244) and 705 (Mel-Juso_PLX4032) unique phosphopeptides from 164, 160, 373 and 386 proteins, respectively (Fig. 1B and 1C). We performed Gene Ontology Functional Annotation analyses, and found that the identified proteins were enriched for cytoskeleton, cell-cell adherens junction and focal adhesion, all of which are known to be linked to p-Tyr signaling (Fig. 1D). These proteins were also enriched for biological processes including transmembrane RTK activity, ephrin receptor binding, non-membrane spanning tyrosine kinase activity and molecular functions including VEGFR signaling pathway and ephrin receptor signaling pathway (Fig. 1D). Consistent with these findings, protein-protein interaction-based subcellular localization prediction analyses revealed that the identified p-Tyr proteins have a higher probability score for plasma membrane or cytoskeleton localization, compared to intracellular organelles including the endosome, Golgi, lysosome and peroxisome (Fig. 1E).

Next we performed quantitative assessment of the data by generating unsupervised hierarchical clustering based on the change of the phosphorylation level of specific Tyr residues. The resulting heatmap showed that these site-specific phosphorylation alterations could segregate the four datasets (Fig. 2A). Specifically, both PLX4032 and AZD6244 treatment led to a profound remodeling of the Tyr phosphoproteome in A375 cells. On the other hand, treatment of Mel-Juso cells with AZD6244, but not PLX4032, reshaped RTK-mediating signaling. These results are consistent with a model where robust blockade of the MAPK pathway is required for the adaptive activation of p-Tyr-mediated signaling (Fig. S1D). From the A375_AZD6244 dataset, we identified 22 and 29 p-Tyr proteins that were up- and down-regulated (by more than 2-fold) after AZ6244 treatment, respectively (Fig. S2A, left). GO analyses of the up-regulated phosphorylated proteins revealed that they are enriched with the transmembrane RTK signaling pathway (e.g., IRS2, ERBB3, STAT3, PTPN11) (Fig. S2A, right). We performed similar analyses on the upregulated phosphoproteins from the PLX4032-treated A375 cells, and found that many of these proteins were also linked to RTK signaling (e.g., IRS2, IL6ST, JAK1, STAT3) (Fig. S2B). From the GO analyses, we observed JAK-STAT pathway was enriched for the upregulated p-Tyr proteins in both A375_PLX4032 and A375_AZD6244 datasets. For example, marked induction of STAT3 Tyr705 phosphorylation was identified by MS in A375 cells after the treatment with PLX4032 or AZD6244 (Fig. S2C, S2D). Activation of JAK/STAT3 signaling is an important adaptive resistance mechanism to EGFR tyrosine kinase inhibitors (TKI) in non–small cell lung cancer (NSCLC) (18). Our data suggest that JAK/STAT3 could also be a bypass mechanism that is relevant to MAPK-inhibited melanoma cells.

**Fig. 2.**
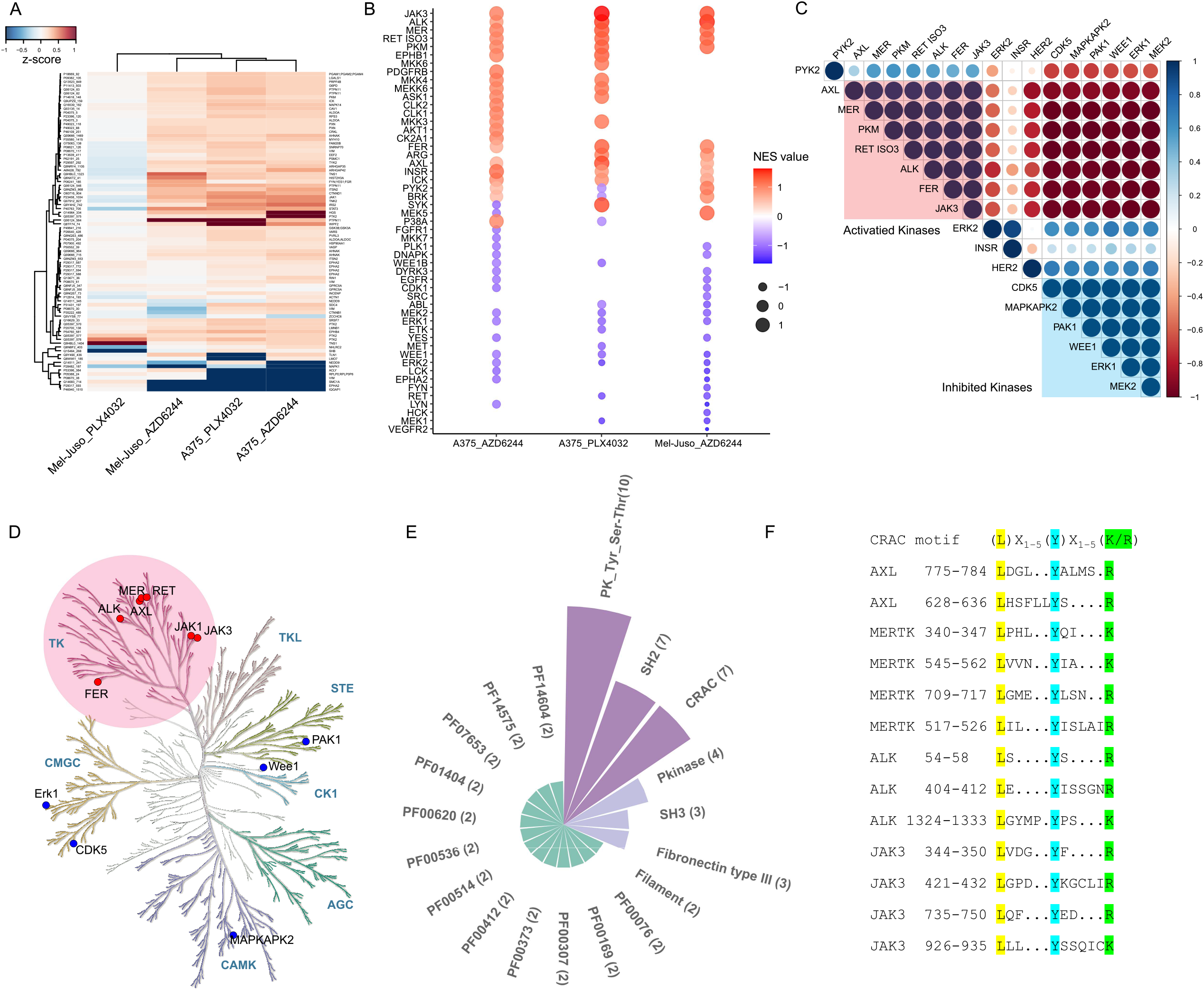
(A) Heatmap showing the unsupervised hierarchical clustering analysis of the phosphorylated peptides commonly quantified across A375_AZD6244, A375_PLX4032, Mel-Juso_AZD6244 and Mel-Juso_PLX4032 experiments. The values represent the z-score for the relative fold change of the ion intensity of the phosphopeptides in each drug treatment condition with respect to DMSO treatment (B) Kinase activity inference results using the scaled phosphopeptide data set for each experiment. Each dot in the plot represents the normalized enrichment score (NES) for a single kinase. (C) The Pearson correlation matrix of the NES of the enriched kinases identified in B. The clusters of activated kinases (red) and inhibited kinases (blue) are highlighted. (D) Phylogenetic kinome tree depicting the protein kinase super families of the enriched kinases in (Fig. 2C). (E) Protein motifs enriched in the upregulated phosphoproteins and activated kinases in A375_AZD6244, A375_PLX4032 and Mel-Juso_AZD6244. (F) Schematic representation of the CRAC consensus sequences in AXL, MERTK, ALK and JAK3.

GO analyses of upregulated p-Tyr proteins enriched in the Mel-Juso_AZD6244 and Mel-Juso_PLX4032 datasets showed that these two compounds induced different p-Tyr patterns. Similar to the A375_AZD6244 dataset, the up-regulated proteins in the Mel-Juso_AZD6244 dataset were also linked to RTK signaling (e.g., the IL-6-JAK/STAT3 signaling pathway) (Fig. S3E). In contrast, the lack of significant functional enrichment of the p-Tyr proteins in PLX4032-treated Mel-Juso cells could be ascribed to the inability of BRAF inhibitors to block MAPK signaling in these NRAS^mut^ cells (Fig. S3F).

### Identification of cholesterol as the master regulator for the adaptive response to MAPK inhibition

The aforementioned quantitative mass spectrometric data showed that MAPK inhibition in two independent cell systems (i.e., A375 and Mel-Juso) both results in a profound remodeling of their p-Tyr proteome. Because of the extensive activation of RTK-mediated signaling, we hypothesized that there could be a conserved “master regulator” that controls the coordinated activation of these RTKs during the adaptive response to MAPK inhibition. Towards this, we performed cross-reference analyses of the three screening experiments (i.e., A375_PLX4032, A375_AZD6244 and Mel-Juso_AZD6244) and extracted the “phosphorylation sites” whose p-Tyr levels were commonly upregulated in all three datasets. We then mapped them to known kinase-substrate relationships to computationally infer the relevant upstream kinase activities (19). To perform the “kinase activation” analysis, lists of kinase-substrate relationships for 348 kinases across a range of kinase families were curated from PhosphoSitePlus (20). Then the “phosphorylation sites” from the MS data were queried against the “kinase-substrate” lists using the GSEA algorithm (21). From the analyses, we found the enriched kinases signatures were similar among A375_AZD6244, A375_PLX4032 and Mel-Juso_PLX4032 (Fig. 2B). The correlation analysis showed that in the drug treated cells, the activity of many kinases, including AXL, MER, RET, ALK, FER and JAK3, etc., were specifically upregulated (activated kinases) (Fig. 2C). Meanwhile, several AZD6244/PLX4032 downstream kinases (MEK2, ERK1 and MAPKAPK2) showed substrate enrichment for the downregulated phosphorylation peptides (inhibited kinases), indicating the drug treatment was still effective when cells were harvested (Fig. 2C). The kinome analysis revealed that the activated kinases were enriched for Tyrosine Kinases (TK) family on the KinMap with different branches (Fig. 2D), indicating AZD6244 or PLX4032 treatment in A375 cells, and PLX4032 treatment in Mel-Juso cells induced the activation of multiple RTKs.

To explore the common features of the activated substrates and tyrosine kinases, we generated a list of 62 proteins, which included the MS-identified commonly upregulated phosphorylated proteins as well as the computationally inferred enriched activated kinases from A375_AZD6244, A375_PLX4032 and Mel-Juso_PLX4032 experiments. Then we queried the list against the Pfam database which is a comprehensive collection of protein families, clans and domains based on the multiple sequence alignments. Intriguingly, the most represented motifs after the analysis were protein kinase domains (PF07714, PF00069), the cholesterol-binding motif known as the <u>**C**</u>holesterol <u>**R**</u>ecognition/interaction <u>**A**</u>mino acid <u>**C**</u>onsensus sequence (CRAC)(22) and the SH3/SH2 domains (PF00017, PF00018) (Fig. 2E). Although generally considered as a structural lipid backbone of the cell membrane, cholesterol can interact with proteins via CRAC to trigger protein dimerization, a key step in the activation of RTKs (22). The existence of CRAC in these activated kinases (Fig. 2F) suggested that altered cholesterol metabolism could be the underlying mechanism for MAPK inhibition-induced RTK activation, and the subsequent adaptive response in melanoma cells.

To directly test this hypothesis, we first used a fluorometric assay to measure the total cholesterol level in A375 cells treated with or without the MEK (AZD6244) and BRAF (PLX4032) inhibitors. We found that both inhibitors (24 hr treatment) resulted in a ~3-fold increase of the cholesterol level (Fig. 3A). It has been well established that the assembly of cholesterol, sphingolipid as well as ganglioside can form a stable membrane sub-compartment called lipid rafts. These nanoscale domains function as a key platform to regulate membrane signaling and trafficking (23). We next labelled A375 cells with the fluorescently-tagged cholera toxin B subunit (FITC-CTXb), which specifically binds to the GM1 ganglioside, and thus, visualizes the cholesterol-rich microdomains (lipid rafts) in the plasma membrane (24). Indeed, confocal microscope analyses showed that AZD6244 treatment dramatically increased the fluorescence intensity of cholesterol-rich microdomains (Fig. 3B). Quantitative analyses showed that the total fluorescence intensity within an individual cell was about 3-fold higher in A375 cells treated with AZD6244 than that in control cells (Fig. S3A, the outlined region was enlarged in Fig. 3B). Using a different microscope modality, namely the total internal reflection fluorescence (TIRF), we confirmed that the plasma membrane localization of cholesterol-rich microclusters was also significantly increased in AZD6244-treated A375 cells (Fig. 3C).

**Fig. 3.**
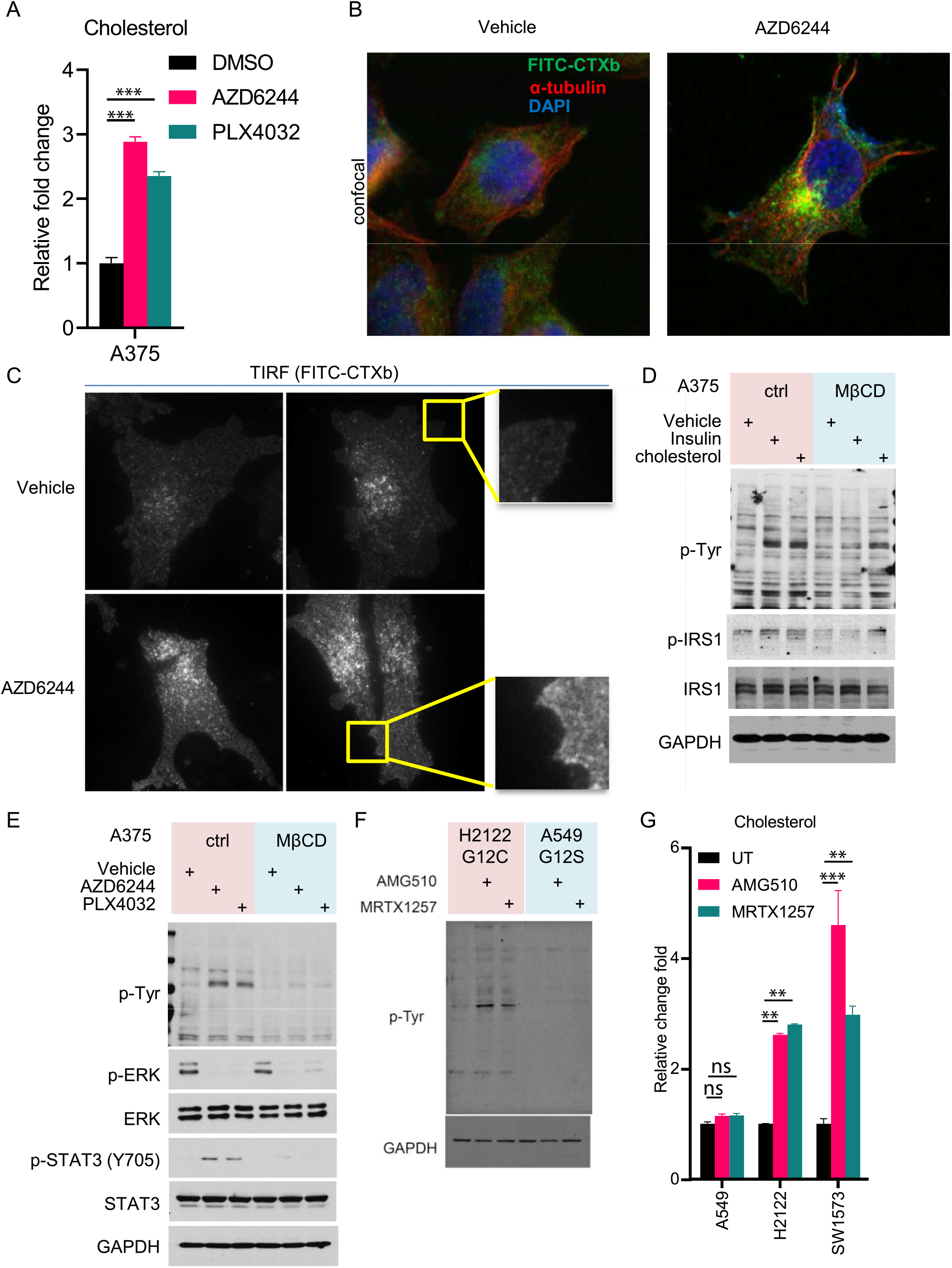
(A) Bar plot of the relative level of cholesterol in A375 cells treated with 1 μM AZD6244 or 1μM PLX4032 for 24 hours. Two-tailed *t* test. ***, p < 0.001. (B) Representative confocal immunofluorescent images of FITC-CTXb, AlexaFluro 594-anti–α-tubulin and DAPI staining of A375 cells under DMSO and AZD6244 treatment (1 μM, 24 h) conditions. The corresponding larger scan was shown in Figure S3A. (C) Representative TIRF images of FITC-CTXb staining of A375 cells under treatment conditions as in B. (D) Immunoblots of cell lysates from serum starved A375 cells pretreated with DMSO or 5 mM MβCD for 1h at 37°C, then stimulated with 200 nM Insulin or 20 mM Cholesterol for 10 min 37°C. GADPH served as a loading control. (E) Immunoblots of cell lysates from A375 cells treated with 1 μM AZD6244 or PLX4032, with or without the presence of 3 mM MβCD for 24 hours. (F) Immunoblots of cell lysates from H2122 and A549 cells treated with 1 μM AMG-510 or MRTX-1257 for 24h at 37°C. (G) Bar plot of the relative level of cholesterol in H2122, SW1573 and A549 cell lysates treated with 1 μM AZD6244 or 1μM PLX4032 for 24 hours. Two-way ANOVA analysis followed by Turkey’s post hoc test. ns, not significant; **, p < 0.05; ***, p < 0.001.

Many growth factor receptors have been shown to be enriched in lipid rafts (25), with their dimerization, and subsequent activation further regulated by cellular cholesterol contents (26, 27). The dramatic increase in both total cholesterol abundances and lipid raft formation in MAPK-inhibited A375 cells prompted us to investigate whether cholesterol could function as the master regulator to control the adaptive remodeling of the p-Tyr proteome. We first confirmed that cholesterol was required for p-Tyr signaling events mediated by various RTKs. As a model system, we found that insulin stimulation in serum-starved A375 cells increased the global p-Tyr level, as well as the phosphorylation of Insulin Receptor Substrate 1 (IRS1), which is a downstream substrate of the Insulin Receptor (IR) (Fig. 3D, Figure 3-source data 1). Similar results were obtained when these cells were treated with cholesterol (Fig. 3D). Pre-treating the cells with methyl-β-cyclodextrin (MβCD), which is a cholesterol-depleting agent(28), blocked insulin-induced global p-Tyr, as well as p-IRS1 signals (Fig. 3D). Pre-complexed MβCD and cholesterol at a 1:2 molar ratio partially rescued p-Tyr inhibition caused by cholesterol depletion (Fig. 3D). We explored the role of cholesterol in RTK activation induced by MAPK inhibitors. In A375 cells, both AZD6244 and PLX4032 treatment increased global p-Tyr and p-STAT3 levels. However, in the presence of MβCD, the induction of global p-Tyr and p-STAT3 was completely abrogated (Fig. 3E, Figure 3-source data 1). Collectively, these data indicate that cholesterol is required for the bypass p-Tyr activation caused by various MAPK inhibitors.

We extended our findings by analyzing the correlation between cholesterol/ lipid abundances and drug sensitivity in a wider panel of melanoma cell lines. The Cancer Cell Line Encyclopedia (CCLE) project used liquid chromatography–mass spectrometry (LC-MS) and profiled more than 225 metabolites in 928 cell lines from more than 20 cancer types (29). We extracted the data from the 40 melanoma cells that harbor the BRAF V600E mutation. Based on their response to AZD6244 and PLX4720 (the tool compound for PLX4032) (data derived from the Cancer Therapeutics Response Portal), we then classified these cell lines into the “sensitive” (ActArea > 3) and “resistant” (ActArea < 3) groups (30). Statistical analyses revealed that the resistant cells have a significantly higher level of cholesterol and other lipid species, compared to the sensitive cell lines (Fig. S3B). Taken together, these results suggest that at least for melanoma cells, increased abundances of cholesterol and the related lipid species could also play a role in determining their innate resistance to MAPK inhibitors.

Besides in melanoma, aberrant activation of MAPK signaling is also found in many other human malignancies (e.g., as a result of KRAS mutation in NSCLC) (31, 32). Marked progresses have been reported for the development of covalent inhibitors targeting the KRAS^G12C^ mutant. Not dissimilar to BRAF/MEK inhibitors for BRAF^mut^ melanoma, long term treatment of KRAS^G12C^ inhibitors, however, also triggers the tumor adaptive response, which is characterized by the reactivation of MEK/ERK under these conditions. We then explored the role of cholesterol in regulating the bypass mechanisms in KRAS^G12C^ tumors. Indeed, treatment of H2122 cells (a KRAS^G12C^ NSCLC cell line) with two chemically distinct, covalent KRAS^G12C^ inhibitors (i.e., AMG 510 and MRTX 1257) resulted a dramatic increase in the global p-Tyr levels (Fig. 3F, Figure 3-source data 1). However, no significant changes of p-Tyr signaling were observed in A549 cells (a KRAS^G12S^ NSCLC cell line) treated with these compounds (Fig. 4A). Accordingly, fluorometric assays showed that there was a 2 to 5-fold increase of the total cholesterol level in H2122 and SW1573 (a KRAS^G12C^ NSCLC cell line) treated with AMG510 or MRTX1257 (Fig. 3G). However, the same treatment in A549 cells had no effect on its cholesterol levels (Fig. 3G). These findings indicate that aberrant cholesterol metabolism might be a conserved mechanism for the adaptive response in tumors with hyperactive MAPK signaling.

**Fig. 4.**
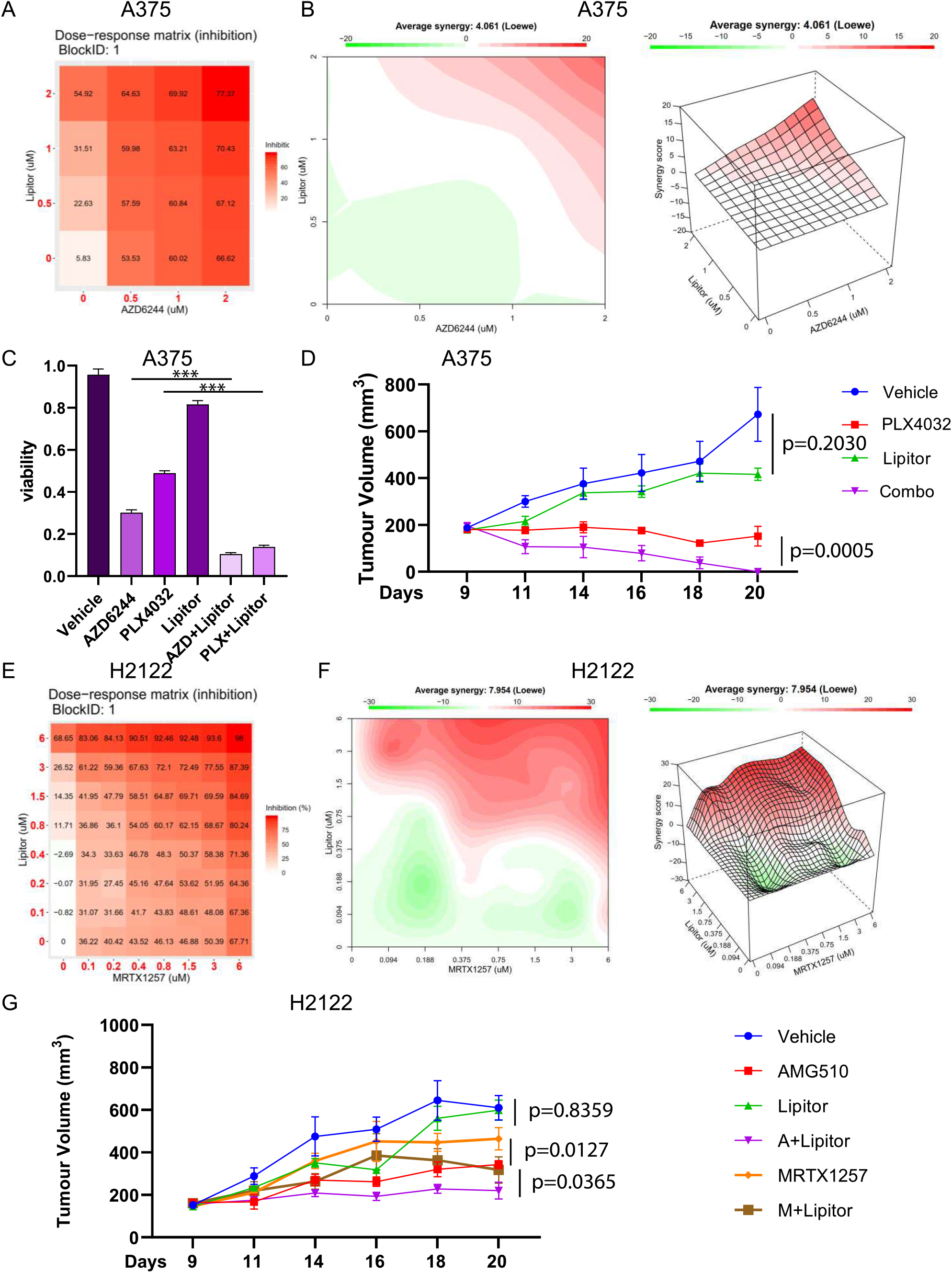
(A) The dose-response matrix of A375 cells treated with AZD6244 and Lipitor by the indicated concentrations for 48 hours at 37°C. The numbers indicate the degree of the cell death (percentages). (B) The synergy plots show that the combination of AZD6244 and Lipitor is synergistic in A375 cells. (C) The viability of A375 cells after the treatment of the indicated compounds for 48 hours at 37°C. The concentration for each compound was 1 μM. Data are means ± SEM with three replicates. One-way ANOVA analysis followed by Turkey’s post hoc test. ***, p < 0.001. (D) Tumor sizes of an A375 xenograft model (*n* = 5 per group). Mice were dosed with 1 mg/kg PLX4032, 1.2 mg/kg Lipitor and the combination for 2 weeks (after 7 days post injection of the tumor cells). Tumor sizes were measured on the indicated days post injection. Two-way ANOVA analysis followed by Turkey’s post hoc test. (E) The dose-response matrix of H2122 cells treated with MRTX1257 and Lipitor using the indicated concentrations for 48 hours at 37°C. The numbers indicate the degree of cell death (percentages). (F) The synergy plots show that the combination of MRTX1257 and Lipitor is synergistic in H2122 cells. (G) Tumor sizes in an H2122 xenograft model (*n* = 5 per group). Mice were dosed with 5 mg/kg AM G-510, 3 mg/kg MRTX-1257, 1.2 mg/kg Lipitor and indicated combinations for 2 weeks after 7 days post injection of the tumor cells. Tumor sizes were measured on the indicated days post injection. Two-way ANOVA analysis followed by Turkey’s post hoc test.

### Synergistic effects between cholesterol-lowering agents and MAPK inhibitors

Cholesterol is being increasingly appreciated for its role in regulating the initiation, progression and metastasis of various human cancers (33, 34). Indeed, large-scale meta-analyses have pointed to decreased cancer incidence in patients that use cholesterol-lowering drugs (e.g., statins) (35, 36). Because of the critical role of cholesterol in regulating the tumor adaptive response to MAPK inhibitors, we sought to test whether cholesterol-lowering drugs could enhance the cytotoxic effects of MAPK inhibitors. Using a CellTiter-Glo assay, we performed a 4 by 4 dose-response validation matrices using Lipitor (Atorvastatin, an FDA-approved, cholesterol-lowering agent), AZD6244 and PLX4032 (Fig. 4A, S4A). The dose-response profiles showed that the combination of Atorvastatin and AZD6244 was highly synergistic in A375 cells, which was evidenced by the positive Loewe score (4.061) (Fig. 4B). We also observed a highly synergistic effect between Atorvastatin and PLX4032, which was associated with a higher Loewe score (14.789) (Fig. S4B). The addition of Atorvastatin to either AZD6244 or PLX4032 treatment also greatly enhanced the cytotoxic effects of these MAPK inhibitors in A375 cells (Fig. 4C, S4C). Next we evaluated the synergistic effects between Atorvastatin and PLX4032 in vivo. In an A375 xenograft model, PLX4032 treatment significantly slowed tumor growth, whereas Atorvastatin alone had no obvious effects. Strikingly, the combination of PLX4032 and Atorvastatin resulted in dramatic regression of the tumors (Fig. 4D).

We also explored the potential synergistic effect between cholesterol-lowering agents and KRAS^G12C^ inhibitors. In this case, MRTX1257 was combined with Atorvastatin in a pairwise, all-versus-all fashion, in 8 × 8 dose-response matrices for three NSCLC cell lines. The results showed that the combination of Atorvastatin and MRTX1257 was highly synergistic in both KRAS^G12C^ lines, i.e., H2122 (Fig. 4E-F) and SW1573 (Fig. S4D-E). However, MRTX1257 had little effect on the proliferation of the KRAS G12S cell line, A549, even at a concentration as high as 5 μM (Fig. S4F). MRTX1257 also did not show any synergistic effect when combined with Atorvastatin (Fig. S4G). Finally, KRAS^G12C^ lung cancer is sensitive to AMG510 or MRTX1257 monotherapy in xenograft experiments, where both KRAS^G12C^ blockers inhibited tumor growth (Fig. 4G). However, the combination of KRAS^G12C^ inhibitors with Atorvastatin was able to achieve a more complete response (Fig. 4G). These findings indicate that besides in melanoma, cholesterol-targeting agents could also overcome the adaptive response in KRAS^G12C^-mutated NSCLC.

## Discussion

Our understanding of how tumor cells develop resistance to targeted kinases inhibitors has been buoyed by rapid progresses in DNA sequencing technologies and has led to some of the first advances in improved therapeutic strategies with more complete and durable responses. Besides the genetic causes, recent studies have shown that resistance can also arise through non-genetic mechanisms. Indeed, activation of alternate routes of kinase pathways is frequently observed following targeted inhibition of oncogenic kinases. An understanding of the molecular underpinnings of these events will be critical for the rationale design of combination approaches to overcome therapeutic resistance. We utilized a quantitative mass spectrometry-based proteomic strategy to comprehensively monitor the dynamic remodeling of the p-Tyr proteome. Indeed, our findings aligned well with the previous studies, with the identification of activated tyrosine kinases (e.g., PDGF, JAK/STAT3, VEGF, c-kit *et al.*) in melanoma cells treated with various MAPK inhibitors (9, 37–40). Among these pathways, the JAK/STAT3 signaling axis has been well documented as a critical mediator of the adaptive tumor response (18, 41–43). However, because of the extensive reprogramming of the p-Tyr signaling network, combining MAPK inhibitors with compounds targeting one or two of these activated kinases is less likely to be sufficient to overcome the adaptive tumor response.

In search for a molecular switch that regulates the coordinated activation of multiple RTKs, we identified a cholesterol binding motif that is commonly shared by many of these tyrosine kinases. Biochemical experiments then confirmed the dramatic accumulation of cholesterol in melanoma cells treated with the MAPK inhibitors. Cholesterol is an important component of the “lipid rafts” and the related structures in the plasma membrane of mammalian cells (44). Lipid rafts was initially studied in the context of protein tyrosine kinase signaling. When bound to their ligands, certain RTKs move into cholesterol-rich lipid microdomains that contain other kinases, adaptors and scaffolds, which become activated to initiate downstream signaling (23, 45, 46). Indeed, we showed that depletion of cholesterol blocks RTK signaling induced by various MAPK inhibitors. Through mining the Cancer Cell Line Encyclopedia (CCLE) database, we also found that cancer cells with higher abundances of cholesterol and the related lipid species are characterized by more innate resistance to MAPK inhibitors. Finally, cholesterol-lowering drugs (e.g., statins) already exhibit beneficial effects by reducing the risk and mortality in multiple cancer types, such as breast, prostate and colorectal cancer (33, 34). Indeed, we found that a profound synergistic effect between Atorvastatin and BRAF/MEK inhibitors in melanoma, as well as between Atorvastatin and a KRAS^G12C^ inhibitor in NSCLC, both in vitro and in vivo.

In addition to RTKs, cellular cholesterol is also involved in regulating the activation of other important signaling molecules. For example, it has been shown that, in order to fulfill its signaling roles, a fraction of H-Ras needs to be localized to the cholesterol-rich lipid rafts in the plasma membrane. The expression of a dominant-negative mutant of caveolin depletes cell surface cholesterol, and thereby inhibits the membrane partition and the subsequent activation of H-Ras (47). Furthermore, lysosomal cholesterol has been shown to activate mTORC1 via the SLC38A9-Niemann-Pick C1 (NPC1) signaling complex (48). These results point to the intriguing possibility that MAPK inhibition-induced cholesterol accumulation could also lead to the compensatory activation of H-Ras and mTORC1 (and perhaps many other signaling molecules), and thereby provide yet other forms of pro-survival signals. Taken together, our results demonstrate cholesterol as the central signaling hub that controls the activation of bypass signaling during MAPK inhibition. The potential regulatory mechanisms that lead to cholesterol accumulation under these conditions warrant further investigation.

In summary, we performed comprehensive characterization of the remodeling of the p-Tyr proteome in melanoma cells treated with various MAPK inhibitors. We showed that the blockade of MAPK signaling leads to the accumulation of cholesterol, which serves as a master regulator to control the coordinated activation of multiple RTKs. Importantly, combination of cholesterol-lowering agents and MAPK inhibitors overcomes the adaptive response in melanoma and NSCLC in cell culture and xenograft models. Because both Atorvastatin and several MAPK inhibitors (e.g., PLX4032) have been already been approved the FDA, this provides an actionable combination therapy that can be rapidly translated and tested in clinical studies.

## Acknowledgments

We are grateful to Dr. Tian Qin for discussions and advices regarding setup of the cholesterol treatment condition. We appreciate the technical assistance and expertise of the Live Cell Imaging Core Facility at UT Southwestern. We also thank the members of the Yu laboratory for their feedback and support.

## Funding

This work was supported, in part, by the NIH (R01GM114160 and R35GM134883) and Welch foundation (I-1800) grants to Y.Y., and the CPRIT training grant RP160157 to X-D.W.

## Materials and Methods

### Cell culture

Human melanoma cell lines A375, Mel-juso and non-small cell lung cancer cell lines H2122, SW1573 and A549 were obtained from the American Type Culture Collection (ATCC, Manassas, VA, USA). All cell lines have been DNA fingerprinted using the PowerPlex 1.2 kit (Promega) and were found to be mycoplasma free using the e-Myco kit (Boca Scientific). Cells were cultured as a monolayer in RPMI-1640 medium supplemented with 10% of fetal bovine serum (Invitrogen). Metabolic Labeling/SILAC cell culture was performed as described previously(12). Cells were maintained in an incubator with a humidified atmosphere of 5% CO2 at 37°C.

### Antibodies and reagents

Phospho-p44/42 MAPK (Erk1/2) (Thr202/Tyr204) (#9101), ERK, Phospho-Stat3 (Tyr705) (#9145), STAT3, Phospho-p90RSK, Phospho-S6 Ribosomal Protein (Ser235/236), p-MEK, Phospho-IRS-1 (Ser302), IRS1, GAPDH antibodies and the bead-conjugated rabbit anti-phosphotyrosine (P-Tyr-1000) were purchased from Cell signaling technology. AZD6244, PLX4032 were purchased from Selleck. AMG-510, MRTX-1257 were purchased from MedChemExpress (MCE). Atorvastatin (calcium salt) (Item No. 10493) was from Cayman chemical. Insulin, MβCD and FITC-CTXb were from Sigma.

### SILAC cell culture

A375 and Mel-Juso cells were grown in light ([^12^C_6_ ^14^N_2_]Lys, [^12^C_6_ ^14^N_4_]Arg) and heavy ([^13^C_6_ ^15^N_2_]Lys, [^13^C_6_ ^15^N_4_]Arg) RPMI (Cambridge Isotope Labs), respectively. Both light and heavy RPMI were supplemented with 10% dialyzed FBS (Invitrogen). For drug treatment, cells cultured in heavy media were treated with 1 μM PLX4032 or 1 μM AZD6244 for 48 hours, while the light cells were treated with DMSO for the same durations.

### Sample preparation for mass spectrometric analysis

The heavy and light cells were lysed in urea buffer (8M urea, 20 mM HEPES pH 7.0, 75 mM β-glycerolphosphate, 1 mM sodium vanadate, 1 mM DTT and 1.5 mM EGTA) and the lysates were combined at a 1:1 ratio. Lysates were reduced by adding DTT to a final concentration of 3 mM, followed by incubation at room temperature for 20 min. Cysteines were alkylated by adding iodoacetamide to a final concentration of 50 mM, followed by incubation in the dark for 20 min. The lysates were diluted to a final concentration of 2 M urea by addition of 100 mM NH4OAC (pH 6.8) and were digested overnight with sequencing-grade trypsin (Promega) at a 1:100 (enzyme:substrate) ratio. Digestion was quenched by addition of trifluoroacetic acid to a final concentration of 0.1% and precipitates were removed by centrifugation at 4,000 rpm for 30 min. Peptides were desalted on SepPak C18 columns (Waters) according to manufacturer’s instructions. Phospho-tyrosine peptides were enriched by PTMScan® Phospho-Tyrosine Rabbit mAb (P-Tyr-1000) Kit (CST). Briefly, lyophilized peptides were resuspended in 1.4 mL Immunoaffinity Purification (IAP) buffer and incubated with the antibody-bead slurry for 2 hr at 4°C. Then the beads were washed with 1mM IAP buffer for two times, and 1mL chilled HPLC water for three times. After elution with 0.15% TFA, the p-Tyr modified peptides were desalted/concentrated with home packed C18 StageTips for LC-MS Analysis.

### Mass spectrometry analysis and data processing

The p-Tyr modified peptide samples were analyzed by LC-MS/MS on an Orbitrap Velos Pro mass spectrometer (Thermo, San Jose, CA) using a top-20 CID (collision-induced dissociation) method (49). MS/MS spectra were searched against a composite database of the human IPI protein database (Version 3.60) and its reversed complement using the Sequest algorithm (Ver28) embedded in an in-house-developed software suite. Search parameters allowed for a static modification of 57.02146 Da for Cys and a dynamic modification of phosphorylation (79.96633 Da) on Ser, Thr and Tyr, oxidation (15.99491 Da) on Met, stable isotope (10.00827 Da) and (8.01420 Da) on Arg and Lys, respectively. Search results were filtered to include <1% matches to the reverse data base by the linear discriminator function using parameters including Xcorr, dCN, missed cleavage, charge state (exclude 1+ peptides), mass accuracy, all heavy or light Lys and Arg, peptide length and fraction of ions matched to MS/MS spectra. Phosphorylation site localization was assessed by the Ascore algorithm (50) based on the observation of phosphorylation-specific fragment ions and peptide quantification was performed by using the CoreQuant algorithm (50, 51).

### Immunoblot analysis

For immunoblot analysis, the cells were extracted in lysis buffer (1% SDS, 10 mM HEPES, pH 7.0, 2 mM MgCl_2_, universal nuclease 20 U/ml), and extracts were mixed with the 5 × reducing buffer (60 mM Tris-HCl, pH 6.8, 25% glycerol, 2% SDS, 14.4 mM 2-mercaptoethanol, 0.1% bromophenol blue). Samples were boiled for 5 min and subject to electrophoresis using the standard SDS–PAGE method. Proteins were then transferred to a nitrocellulose membrane (Whatman). The membranes were blocked with a TBST buffer (25 mM Tris-HCl, pH 7.5, 150 mM NaCl, 0.05% Tween 20) containing 3% non-fat dried milk, and probed overnight with primary antibodies (1:1,000 dilution) at 4 °C and for 1 h at room temperature with peroxidase-conjugated secondary antibodies. Blots were developed using enhanced chemiluminescence, exposed on autoradiograph film and developed using standard methods as described(52).

### Immunofluorescence staining

A375 cells were cultured in 35 mm glass bottom dishes (MatTek), incubated with fluorescein-conjugated cholera toxin B subunit (FITC-CTXb; Sigma) at a concentration of 10 μg/ml in D-PBS for 30 min at 37 °C. Cells were then washed with D-PBS and treated with 1 μM AZD6244 in complete RPMI-1640 medium for 24 h.

For confocal fluorescence microscopy analysis, cells were fixed with 4 % PFA, permeabilized with 0.25 % Triton X-100, then stained with appropriate primary antibodies followed by Alexa Fluor antibodies (Life Technologies) as secondary antibodies. Counter staining of cell nuclei was performed using DAPI (Santa Cruz Biotechnology). Immunofluorescence images were acquired with Zeiss LSM880 Airyscan confocal laser scanning microscope (Zeiss) with ×63 glycerol-immersion objective and scanning resolution of 512 × 512 pixels, zoom factor 6.4 for a subset of images. Immunofluorescence intensity were analyzed using ImageJ Software (NIH, version 1.52).

For Total internal reflection fluorescence structured illumination microscopy (TIRF-SIM) analysis, cells were fixed with 4 % PFA, and mounted in ProLong™ Gold Antifade Mountant (Thermo Fisher). Immunofluorescence images were acquired with DeltaVision OMX SR imaging system (Cytiva).

### Cell viability measurement

Cell viability was measured using the CellTiter-Glo assay kit (Promega). Briefly, after drug treatment, room temperature CellTiter-Glo reagent was added 1:1 to each well and the plates were incubated at room temperature for 2 min. Luminescence was measured with the Synergy HT Multi-Detection Microplate Reader (BioTek) and was normalized against control cells treated with DMSO.

### Cholesterol extraction and measurement

Three to five million cells were homogenized into 200 μL chloroform-methanol (v /v = 2:1), centrifuged for 10 min at 12,000 g at 4 °C. The organic phase was transferred to a clean tube and vacuum dried. The abundance of cholesterol was determined by the Cholesterol Fluorometric Assay Kit (Cayman). Briefly, the dried organic cell extraction was reconstituted in 200 μL assay buffer. In a typical experiment, 50 μL of the sample and a series dilution of the standard were added into each well of a 96 well plate. Fifty microliters of the assay cocktail (4.745 mL assay buffer, 150 μL cholesterol detector, 50 μL HRP, 50 μL cholesterol oxidase, and 5 μL cholesterol esterase) was added in the well and mixed. After 30 min incubation at 37 °C in the dark, the fluorescence signal was determined by a plate reader using excitation wavelengths between 530-540 nm and emission wavelengths between 585-595 nm. The relative abundance of cholesterol was normalized to the DMSO treatment group.

### In vivo drug treatment experiments

PLX4032, AMG-510, and MRTX-1257 were dissolved in 2% DMSO + 30% PEG 300 + 5% Tween 80 + ddH2O. Tumors were engrafted in NSG (NOD-SCID) mice (The Jackson Laboratory) by subcutaneous injection of 3 × 10^5^ A375 or H2122 cells in RPMI-1640 medium supplemented with 50% Matrigel (BD Biosciences, cat. no. 354234). Seven days after the injection, animals were assigned randomly to control and various treatment groups (n = 5 for each group). Tumor bearing mice were administered by oral garvage: (1) Vehicle, 2% DMSO + 30% PEG 300 + 5% Tween 80 + ddH2O; (2) PLX4032, 1 mg/kg body weight/day; (3) Lipitor, 1.2 mg/ kg body weight/day; (4) AMG-510, 5 mg /kg body weight/day; (5) MRTX-1257, 3 mg/ kg body weight/day. The mice were treated every other day. Tumors were measured with an external caliper, and the volume was calculated as (4π/3) × (width/2)^2^ × (length/2).

### Statistical analysis

Quantifications were analyzed by software GraphPad prism version 8 and presented as median with SEM. Statistical significance was determined by t test, one-way or two-way analysis of variance (ANOVA) with Tukey’s multiple comparison post hoc test for comparisons involving more than two groups. In many experiments, the mean values of the control groups were set to 1, and all other values were expressed as fold changes compared with the respective controls. *P* < 0.05 was considered significant, and significance was represented as per the GraphPad prism software. For in-silico analyses, the cutoffs, thresholds, and other statistical measurements are mentioned in the individual sections.

### In-silico analyses

Gene Ontology analysis was performed with DAVID and the selection of Gene Ontology Terms for visualization was curated manually. The most relevant pathways were identified by Ingenuity pathway analysis. A protein interaction network with the protein subcellular localization analysis was generated using the STRING database (Version 11.0). The interaction network was visualized with Cytoscape (3.8.1). Benchmarking substrate-based kinase activity inference was analyzed in gene set enrichment analysis (GSEA). The correlation coefficient analyses, motif enrichment analyses were performed in R 3.6.1. Synergy scoring model analysis was built with the synergyfinder package (2.2.4).in R 3.6.1.

### Data availability

The mass spectrometry data have been deposited to the ProteomeXchange Consortium (https://www.ebi.ac.uk/pride/archive/) via the PRIDE partner repository with the dataset identifiers: PXD021877. Computer code and all the other data supporting the findings of this study are available from the corresponding author upon request.

## Figure Legends

**Fig. S1.**
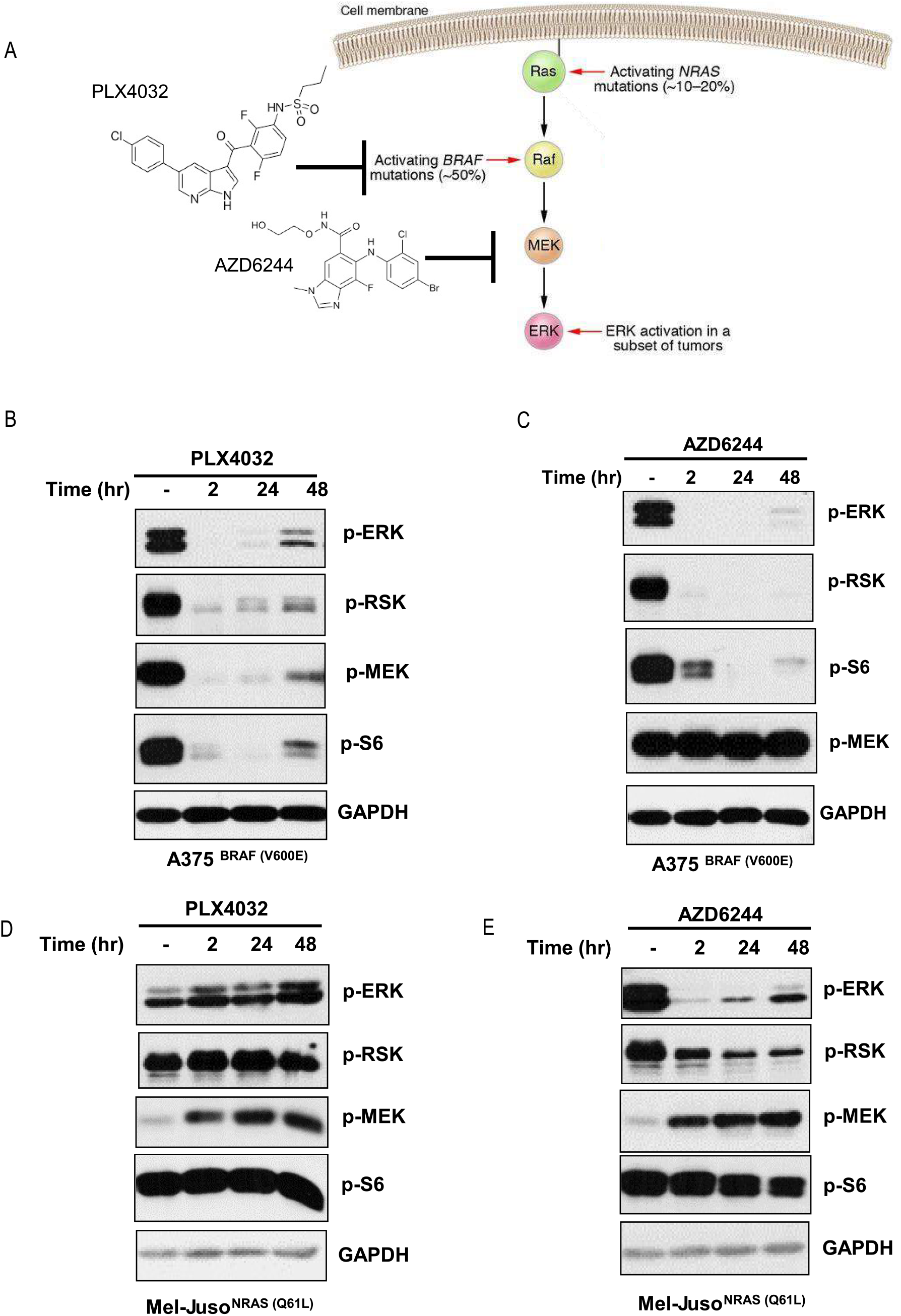
(A) The schematic representation of the RAS-RAF-MEK-ERK signaling pathway. The structures of PLX4032 (a BRAF V600E inhibitor) and AZD6244 (a MEK inhibitor) are shown. (B-C) Immunoblots of cell lysates from A375 cells treated with 1 μM PLX4032 (B) or 1 μM AZD6244 (C) for the indicated times. GADPH served as a loading control. (D-E) Immunoblots of cell lysates from Mel-Juso cells treated with 1 μM PLX4032 (D) or 1 μM AZD6244 (E) for the indicated times. GADPH served as a loading control.

**Fig. S2.**
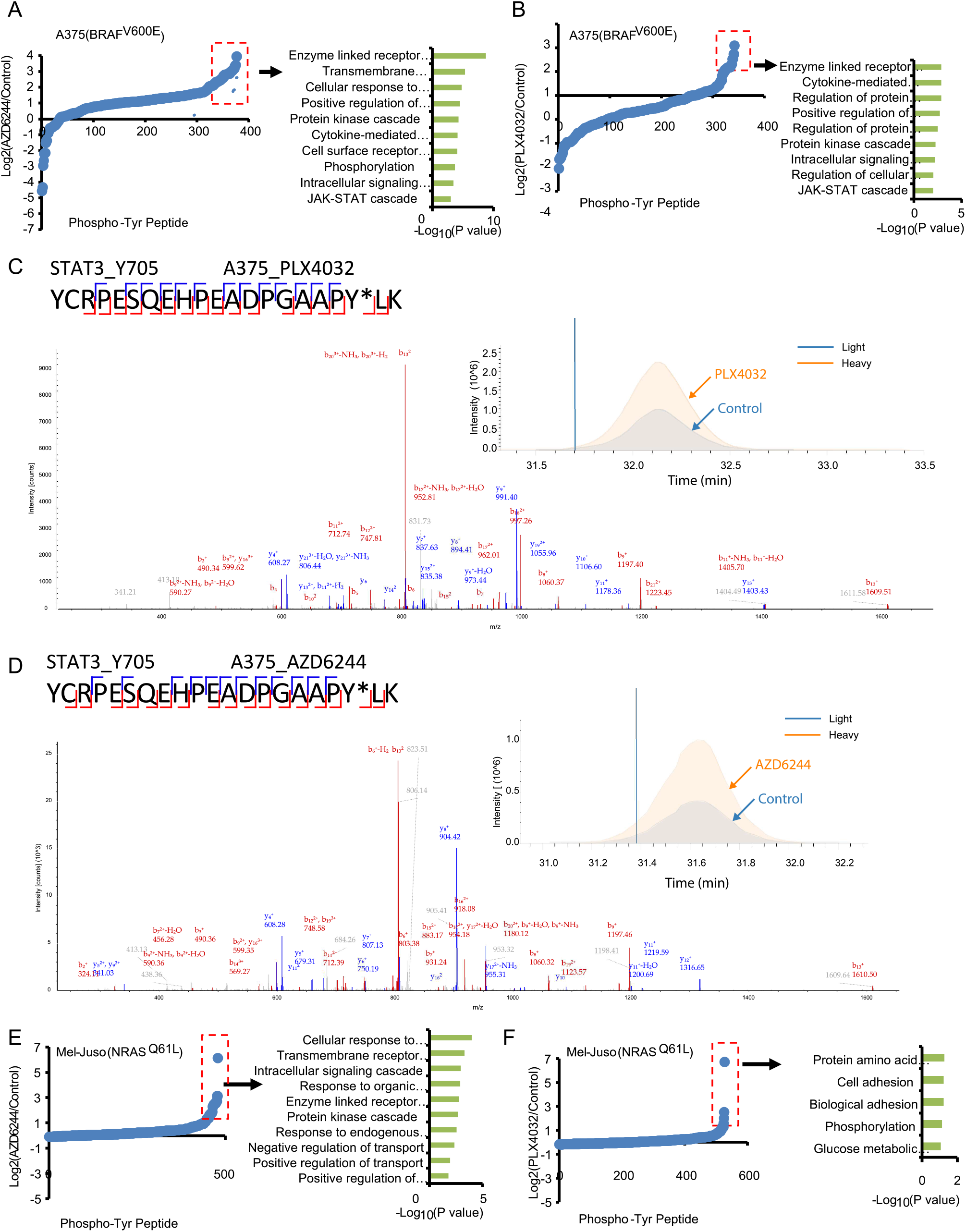
(A-B, E-F) The ratio distribution of phosphopeptides identified and the GO analyses for the upregulated phosphoproteins in each screen: A375_AZD6244 (A), A375_PLX4032 (B), Mel-Juso_AZD6244 (E) and Mel-Juso_PLX4032 (F). Phosphopeptides identified in each screen were extracted and their corresponding treatment/control ratios were plotted on a Log2 scale. Note that most of the phosphopeptides have a ratio of 1:1 between the light and heavy populations and hence have a value close to 0 on a Log2 axis. Proteins with up-regulated phosphorylation (Log2(treatment/control) > 1) for GO analyses in each screen are highlighted in the red box. The pathways enriched in the up-regulated phospho-proteins were plotted on the right. (C-D) The representative tandem mass spectrometry (MS/MS) spectra assignment of the phosphopeptide YCRPESQEHPEADPGAAPY*LK from STAT3 (Y705) identified in A375_PLX4032 (C) and A375_AZD6244 (D). The control and compound-treated profiles are highlight in grey and orange, respectively.

**Fig. S3.**
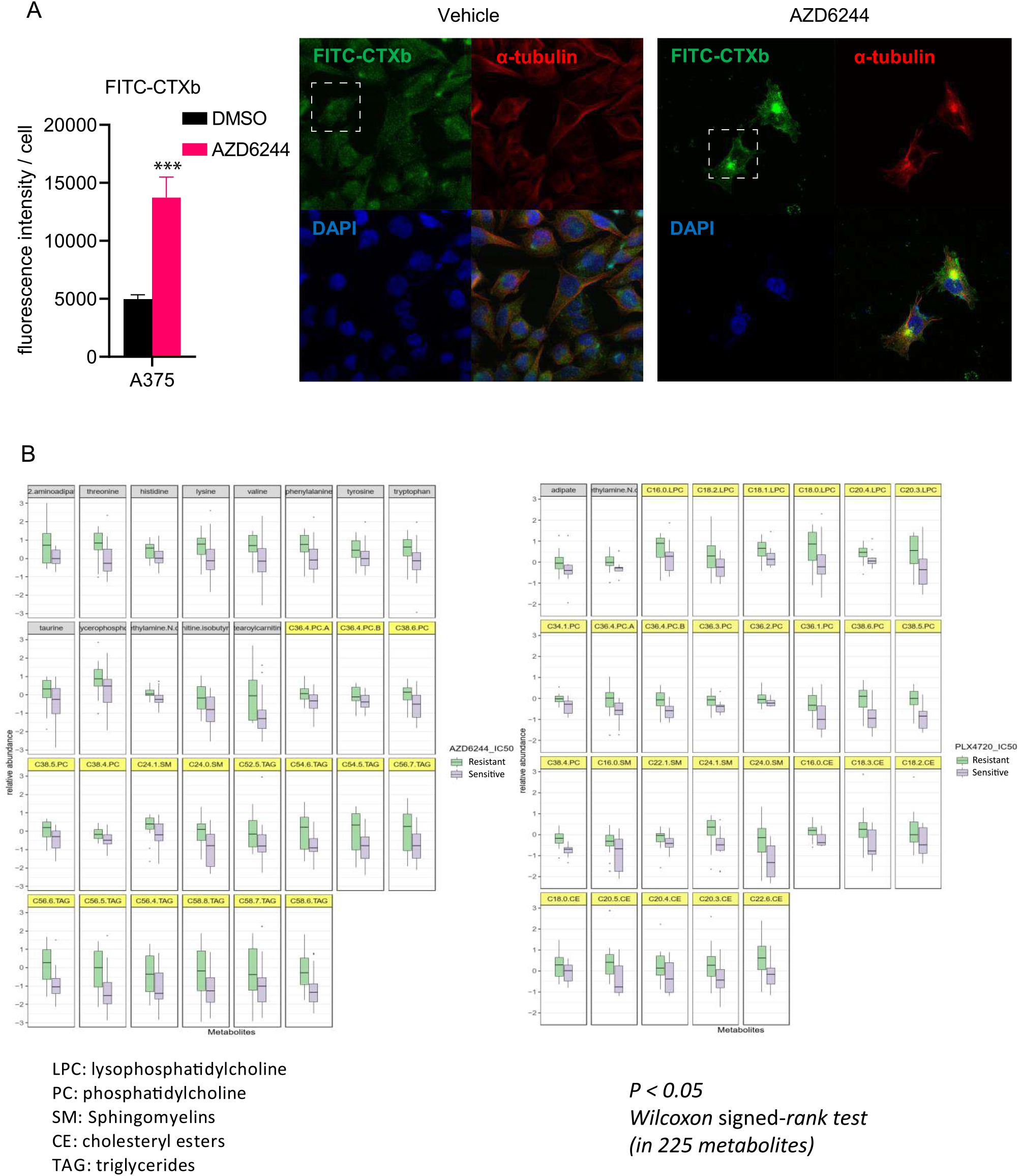
(A) The MFI (mean fluorescence intensity) of FITC-CTXb in A375 cells treated as in Fig. 3B. Two-tailed *t* test. ***, p < 0.001. (B) The metabolite panel of 40 melanoma cells that harbor the BRAF V600E mutation. The cells are classified as “sensitive” (purple) or “resistant” (green) groups in response to AZD6244 and PLX4720 treatment. Data were derived from the Cancer Cell Line Encyclopedia (CCLE) project and the Cancer Therapeutics Response Portal (CTRP). The lipid panel is highlighted in yellow. Wilcoxon signed-rank test, p < 0.05.

**Fig. S4.**
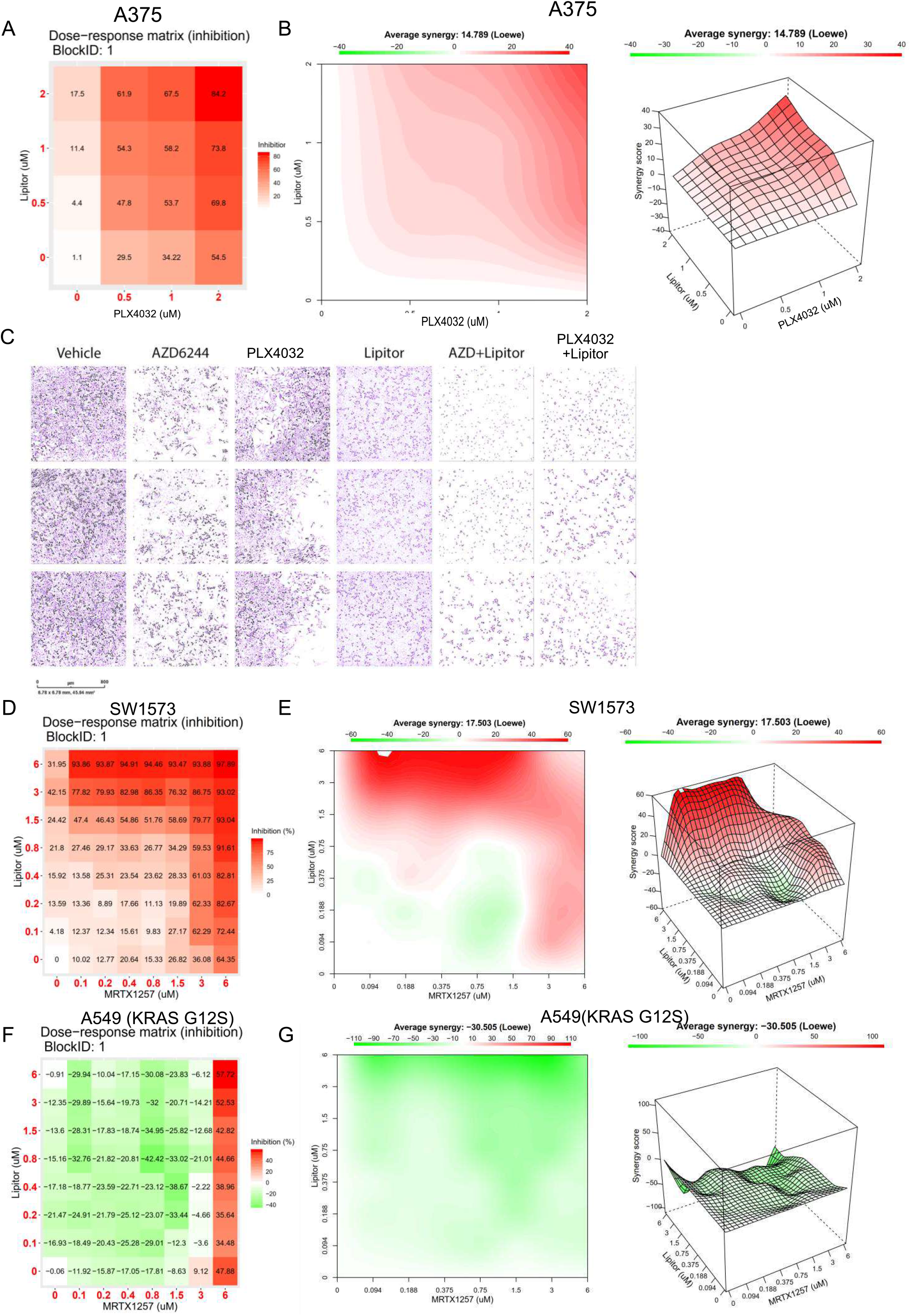
(A) The dose-response matrix of A375 cells treated with PLX4032 and Lipitor using the indicated concentrations for 48 hours at 37°C. The numbers indicate the degree of cell death (percentages). (B) The synergy plots show that the combination of PLX4032 and Lipitor is synergistic in A375 cells. (C) The colony formation assay of A375 cells after the indicated treatment conditions. (D) and (F) The dose-response matrix of SW1573 cells (D) and A549 cells (F) treated with MRTX1257 and Lipitor using the indicated concentrations for 48 hours at 37°C. The numbers indicate the degree of cell death (percentages). (E) and (G) The synergy plots show that the combination of MRTX1257 and Lipitor is synergistic in SW1573 (E) and A549 (G) cells.

**Figure 3-source data 1.**
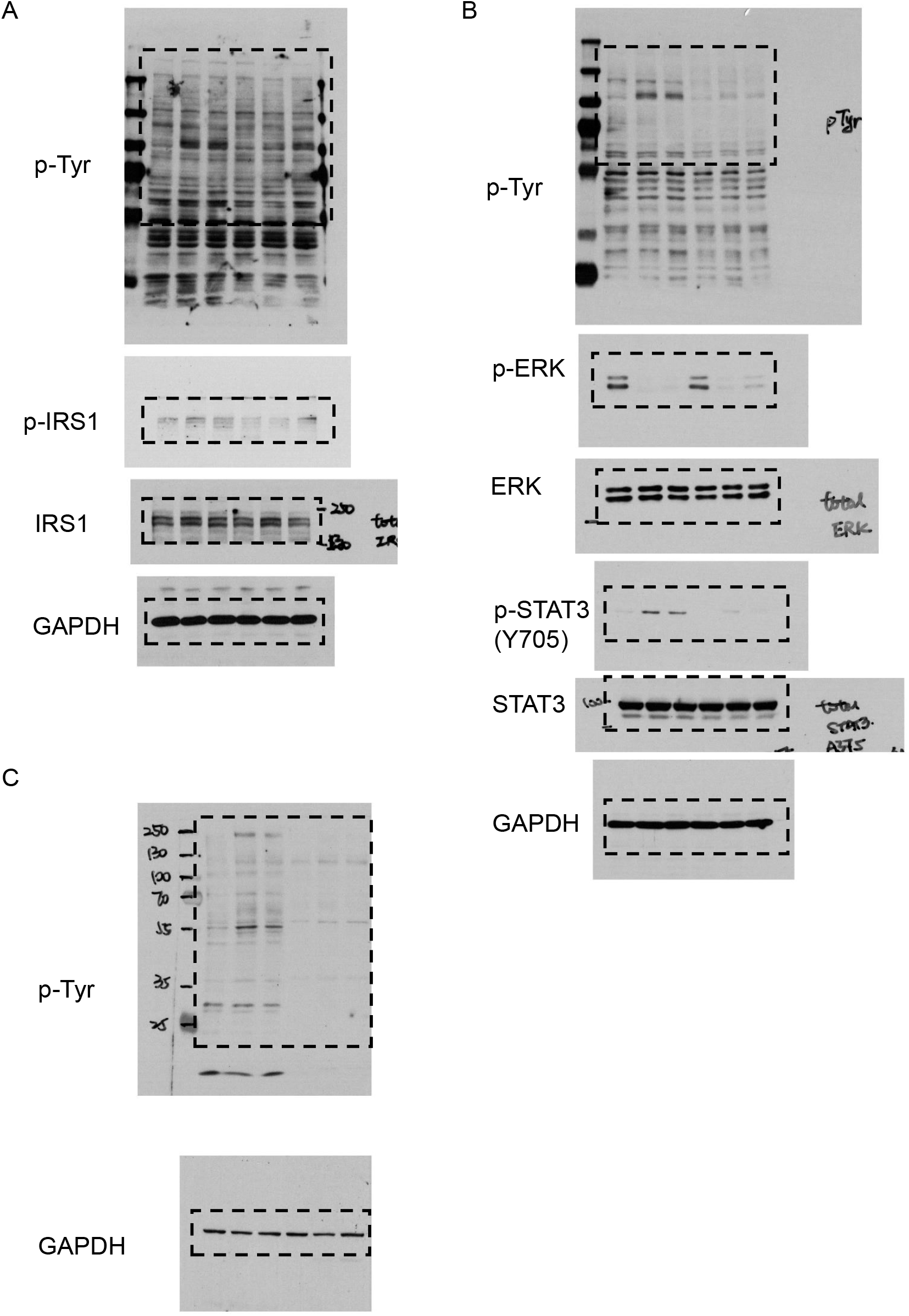
(A) The uncropped blots of Fig. 3D (B). The uncropped blots of Fig. 3E (C). The uncropped blots of Fig. 3F

**Figure S1-source data 1.**
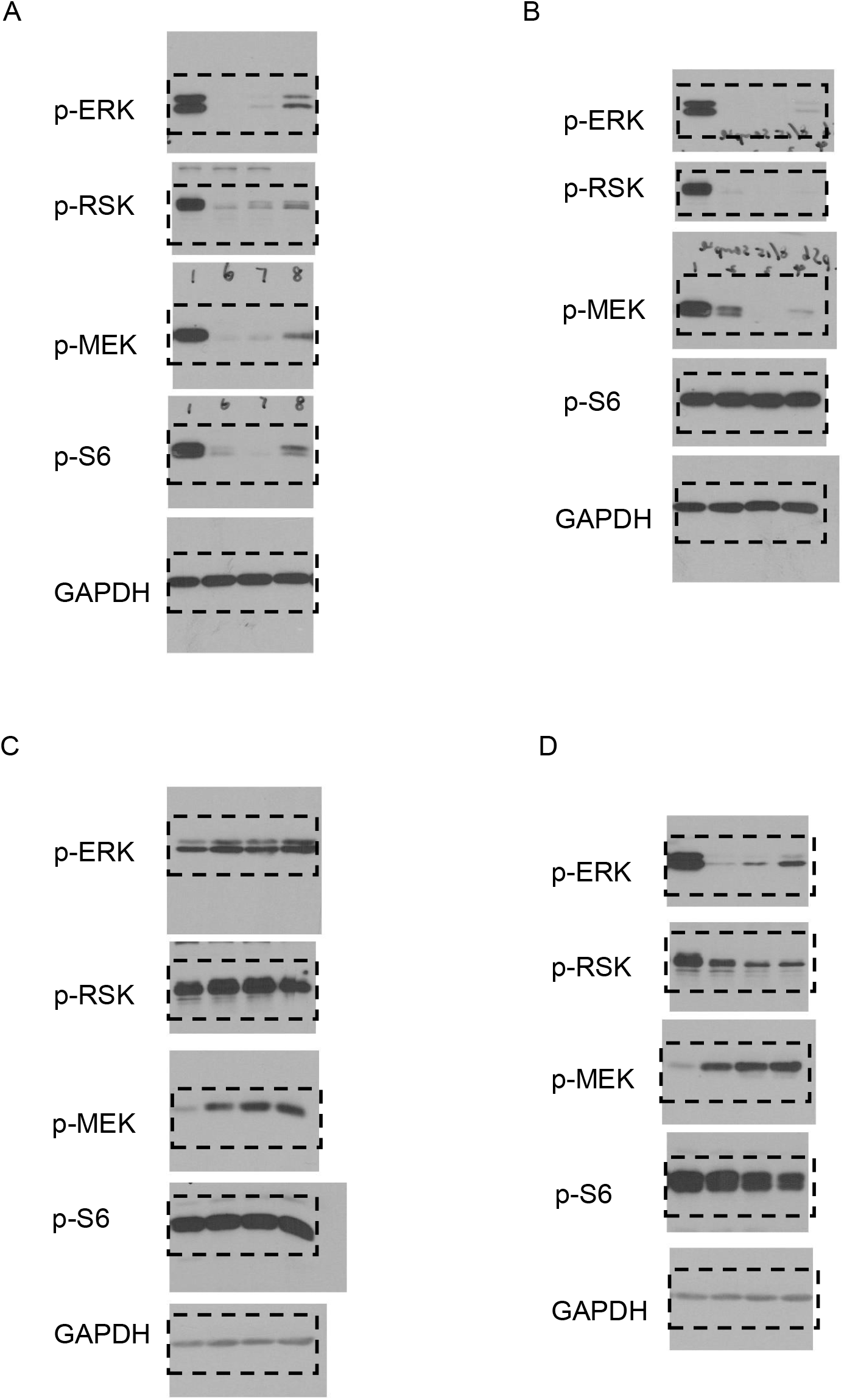
(A). The uncropped blots of Fig. S1B (B). The uncropped blots of Fig. S1C (C). The uncropped blots of Fig. S1D (D). The uncropped blots of Fig. S1E

## Notes

http://proteomecentral.proteomexchange.org/cgi/GetDataset?ID=PXD021877

### Competing Interest Statement

Y.Y. has received research funding from Pfizer.

